# Visualizing Functional Network Connectivity Differences Using an Explainable Machine-learning Method

**DOI:** 10.1101/2024.12.18.629283

**Authors:** Mohammad S. E. Sendi, Vaibhavi S. Itkyal, Sabrina J. Edwards-Swart, Ji Ye Chun, Daniel H. Mathalon, Judith M. Ford, Adrian Preda, Theo G.M. van Erp, Godfrey D. Pearlson, Jessica A. Turner, Vince D. Calhoun

**Affiliations:** Wallace H. Coulter Department of Biomedical Engineering, Georgia Institute of Technology and Emory University, Atlanta, Georgia; Department of Electrical and Computer Engineering, Georgia Institute of Technology, Atlanta, Georgia; McLean Hospital and Harvard Medical School, Boston, MA, USA; Tri-institutional Center for Translational Research in Neuroimaging and Data Science: Georgia State University, Georgia Institute of Technology, Emory University Atlanta, Georgia; Department of Neuroscience, Emory University, Atlanta, Georgia; Department of Computer Science, Georgia State University, Atlanta, Georgia; Department of Psychiatry, Weill Institute of Neurosciences, University of California, San Francisco, CA, United States; Mental Health Service, Veterans Affairs San Francisco Healthcare System, San Francisco, CA, United States; Department of Psychiatry and Human Behavior, University of California, Irvine, Irvine, CA, United States; Department of Psychiatry, School of Medicine, Yale University, New Haven, CT, United States; Department of Psychology, Neuroscience Institute, Georgia State University, Atlanta, GA, United States; Department of Psychiatry and Behavioral Health, College of Medicine, The Ohio State University, Columbus, United States

**Author notes:** (Mohammad S. E. Sendi), (Vaibhavi Itkyal).

## Abstract

Functional network connectivity (FNC) estimated from resting-state functional magnetic resonance imaging showed great information about the neural mechanism in different brain disorders. But previous research has mainly focused on standard statistical learning approaches to find FNC features separating patients from control. Although machine learning approaches provide better models separating controls from patients, it is not straightforward for these approaches to provide intuition on the model and the underlying neural process of each disorder. Explainable machine learning offers a solution to this problem by applying machine learning to understand the neural process behind brain disorders. In this study, we introduce a novel framework leveraging SHapley Additive exPlanations (SHAP) to identify crucial Functional Network Connectivity (FNC) features distinguishing between two distinct population classes. Initially, we validate our approach using synthetic data. Subsequently, applying our framework, we ascertain FNC biomarkers distinguishing between, controls and schizophrenia patients with accuracy of 81.04% as well as middle aged adults and old aged adults with accuracy 71.38%, respectively, employing Random Forest (RF), XGBoost, and CATBoost models. Our analysis underscores the pivotal role of the cognitive control network (CCN), subcortical network (SCN), and somatomotor network (SMN) in discerning individuals with schizophrenia from controls. In addition, our platform found CCN and SCN as the most important networks separating young adults from older.

## 1. Introduction

In recent years, functional connectivity and it’s analog, functional network connectivity (FNC) obtained from resting-state functional magnetic resonance imaging (fMRI) time series have revealed a great deal of knowledge about these brain dysconnectivity in schizophrenia and discriminating these patients from healthy subjects (Lynall *et al* 2010, Skåtun *et al* 2017). But having a limited number of samples with highly dimensional FNC features makes the diagnosis challenging. To overcome this problem, machine learning-based classifications are used to classify one subject class from another based on the FNC data (SZ: Arbabshirani *et al* 2013, 2014, Cao *et al* 2018, Yan *et al* 2017, Jianlong Zhao, Dongmei Zhi, Weizheng Yan, Vince D. Calhoun 2019, Mendoza, R. Cabral-Calderin, Y., Domînguez, M., Garcia, A., Borrego, M., Caballero, A. 2011; Aging: Meunier, D. et al 2009, Zhai, J. et al 2019, Sala-Llonch. Et al 2014; Other: He *et al* 2017a, He *et al* 2017b). However, these approaches do not naturally provide insights into the underlying mechanism of brain FNC affected by the disease. Linear and logistic regression models can more easily explain underlying decision mechanisms taken by the model in a prediction or classification problem. However, we need to sacrifice the model performance in terms of classification accuracy. Using more complex models like random forests, decision trees, and gradient boosted trees or deep learning models, we usually improve the best classification accuracy. However, because of the nonlinear structure of the model, it can be challenging to interpret the model. Therefore, there is always a trade-off between model interpretability and model accuracy in the classification task.

Recent developments in explainable machine learning open a new avenue in excavating the difference between FNC of healthy brain from disease group (Böhle *et al* 2019, Lundberg and Lee 2017, Lundberg *et al* 2018, 2020). In this paper, we developed a framework which can quantify the difference between the whole brain FNC of patients from controls. In more detail, we trained different models differentiating two classes (SZ vs HC or aging vs control) of data. Next, we used the SHapley Additive exPlanations (SHAP) approach (Lundberg and Lee 2017, Lundberg *et al* 2018) as a feature learning method to explain the model and find a subset of the most important features that contribute to this classification.

## 2. Methods

The framework that we have proposed is shown in **Figure 1**. Firstly, we preprocess the rs-fMRI data and extract independent components using a whole brain automated Neuromark pipeline (Du, Y. et al 2020) for the whole brain. Next, we estimate the whole-brain FNC for each participant. Next, the participants are classified into two classes (for instance-patient vs. control). Finally, we use the SHapley Additive exPlanations (SHAP) method, which extracts a subset of features that have the largest contribution to the classification between two classes. Following subsections describe each step in more detail.

**Figure 1:**
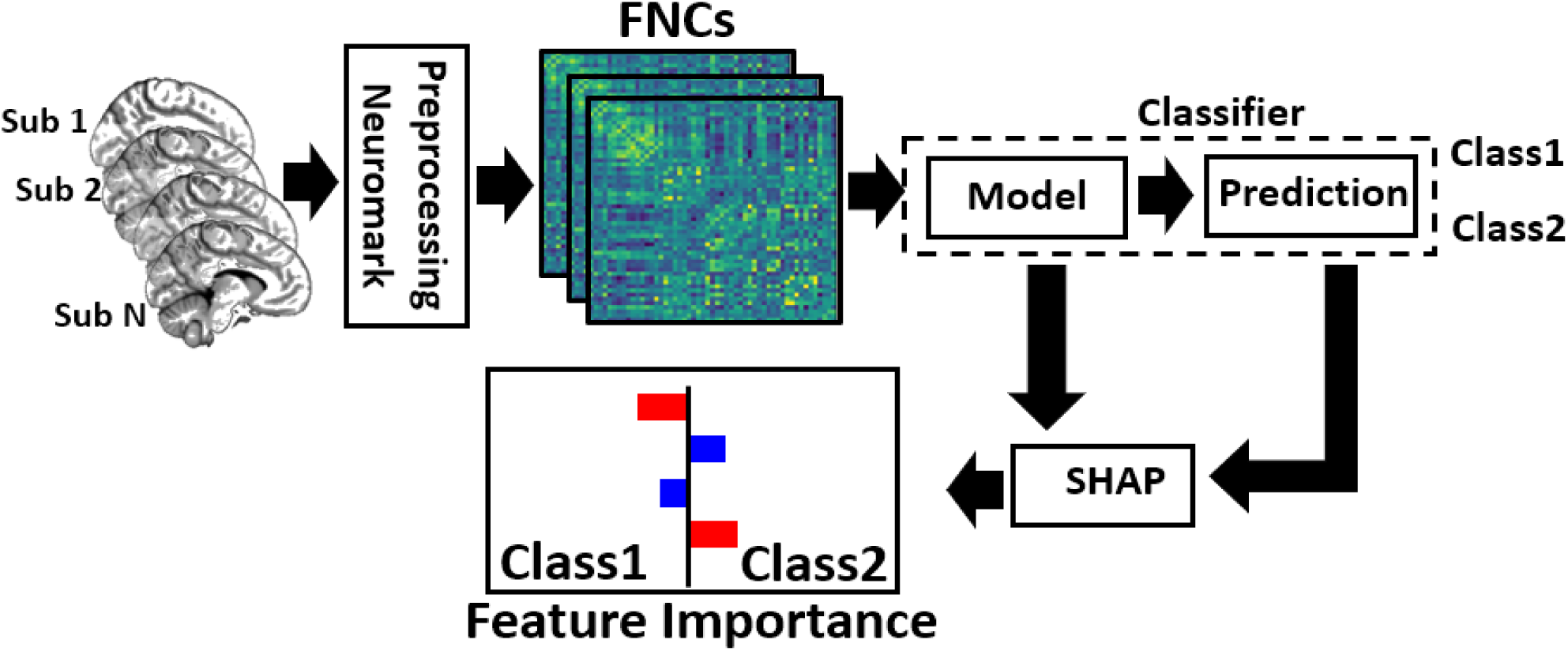
Illustration of our proposed framework. In the first step we processed the fMRI data for all the subjects. Then, using Neuromark, we found the independent components. Next, we calculated the FNC of each subject. These FNC features were fed to the classifier to differentiate between middle adult and old subjects (based on their age). Then, the model and the classifier output are fed to the SHAP explanator which finds the most important features which contribute to the classification between the two classes.

### 2.1 Pre-processing and intrinsic component extraction

To preprocess the fMRI data we applied statistical parametric mapping (SPM12, http://www.fil.ion.ucl.ac.uk/spm/) in the MATLAB2019 environment. We performed slice-timing correction on the fMRI data and then applied rigid body motion correction to correct subject head motion. Next, we performed spatial normalization to an echo planar imaging (EPI) template in the standard Montreal Neurological Institute (MNI) space and resample to 3×3×3 mm^3^. Finally, we used a Gaussian kernel with a full width at half maximum (FWHM) = 6 mm to smooth the fMRI images. In the next step, to extract reliable independent components (ICs), we used the Neuromark automated independent component analysis (ICA) pipeline as introduced in (Du *et al* 2020). In the pipeline, we first identified the replicable components by matching group-level spatial maps from two large-sample healthy control datasets. Then, a subset of matched components was identified as meaningful if they exhibit peak activations in gray matter; have low spatial overlap with known vascular, ventricular, motion, and susceptibility artifacts; and have dominant low-frequency fluctuations on their time-courses. Then, we categorized ICNs into seven domains including subcortical network (SCN), auditory network (ADN), sensorimotor network (SMN), visual network (VSN), cognitive control network (CCN), default-mode network (DMN), and cerebellar network (CBN) based on anatomy and prior knowledge. Total, we extracted 53 ICs for the whole brain. Figure 2 showed all seven networks identified by Neuromark. Table 1 showed all 53 ICs used in this study.

**Figure 2:**
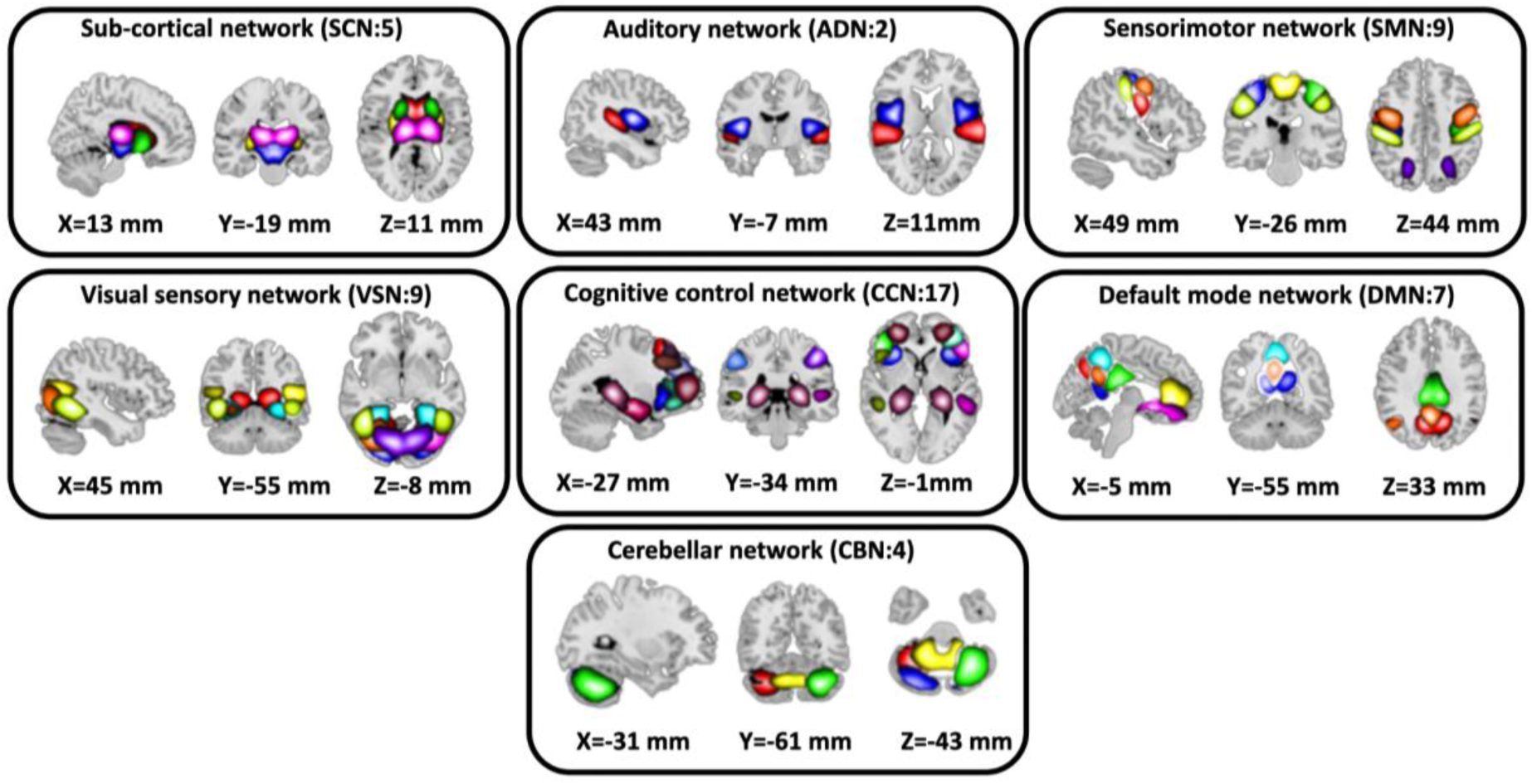
Extracted Independent Components. Neuromark pipeline was used to estimate the 53 ICs and were further put in 7 domains - SCN, AND, SMN, VSN, CCN, DMN and CBN.

**Table 1:**
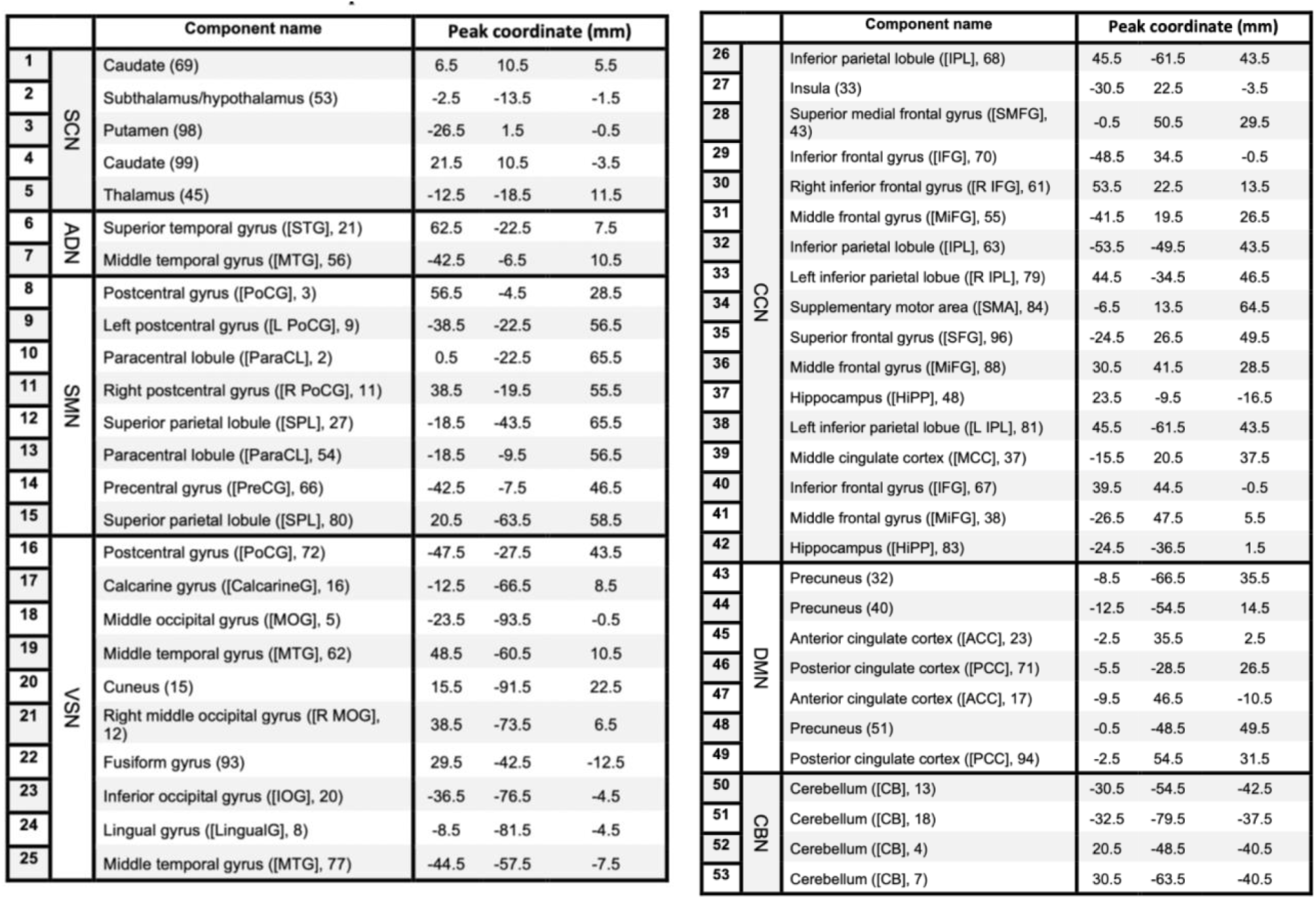
Component labels.

### 2.2 Functional Network Connectivity

To estimate the FNC or the communication strength in the brain, we calculated the Pearson correlation between pairs of ICs in each subject as shown in equation 1

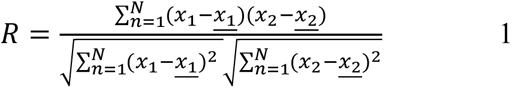

where *x*1 and *x*2 are time course signals and <u>*x*1</u> and <u>*x*2</u> are the mean of *x*1 and *x*2, respectively. It takes values in the interval [− 1, 1] and measures the strength of the linear relationship between *x*1 and *x*2. Each FNC is a 53×53 matrix with 53 ICs. Thus, we calculated 1378 connectivity features for each sample. The FNC in the synthetic dataset is shown in **Figure 4**.

**Figure 3:**
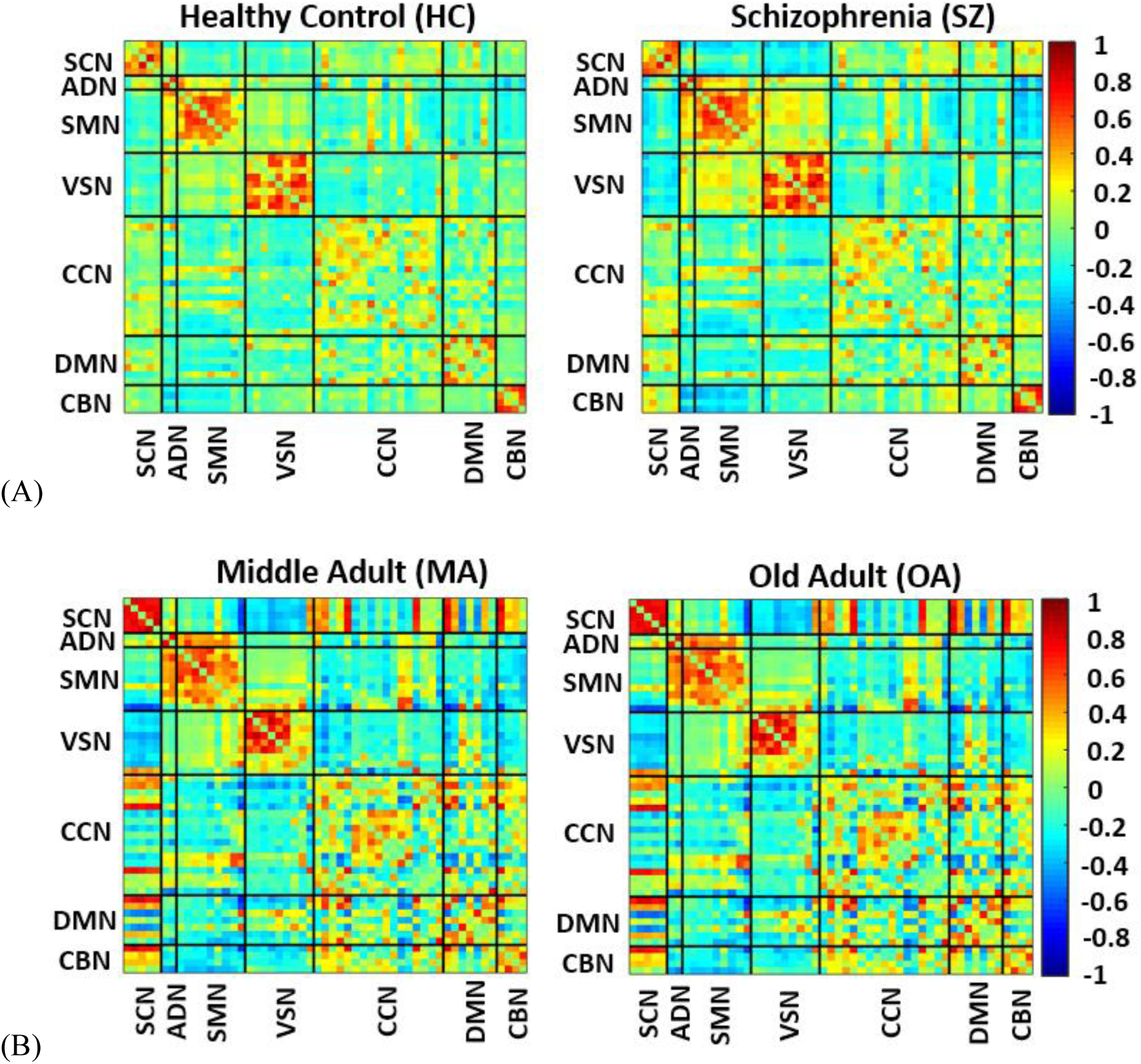
Functional network connectivity (FNC) depicts the correction between pairs of independent components. A) The average FNC across all middle age subjects (<63) and across all old age subjects (>63) in the UK Biobank. B) The average FNC across SZ and HC in the FBIRN dataset. SCN: subcortical network, ADN: Auditory network, SMN: Sensorimotor network, VSN: Visual sensory network, CCN: Cognitive control network, DMN: Default mode network, CBN: Cerebellar networ

**Figure 4:**
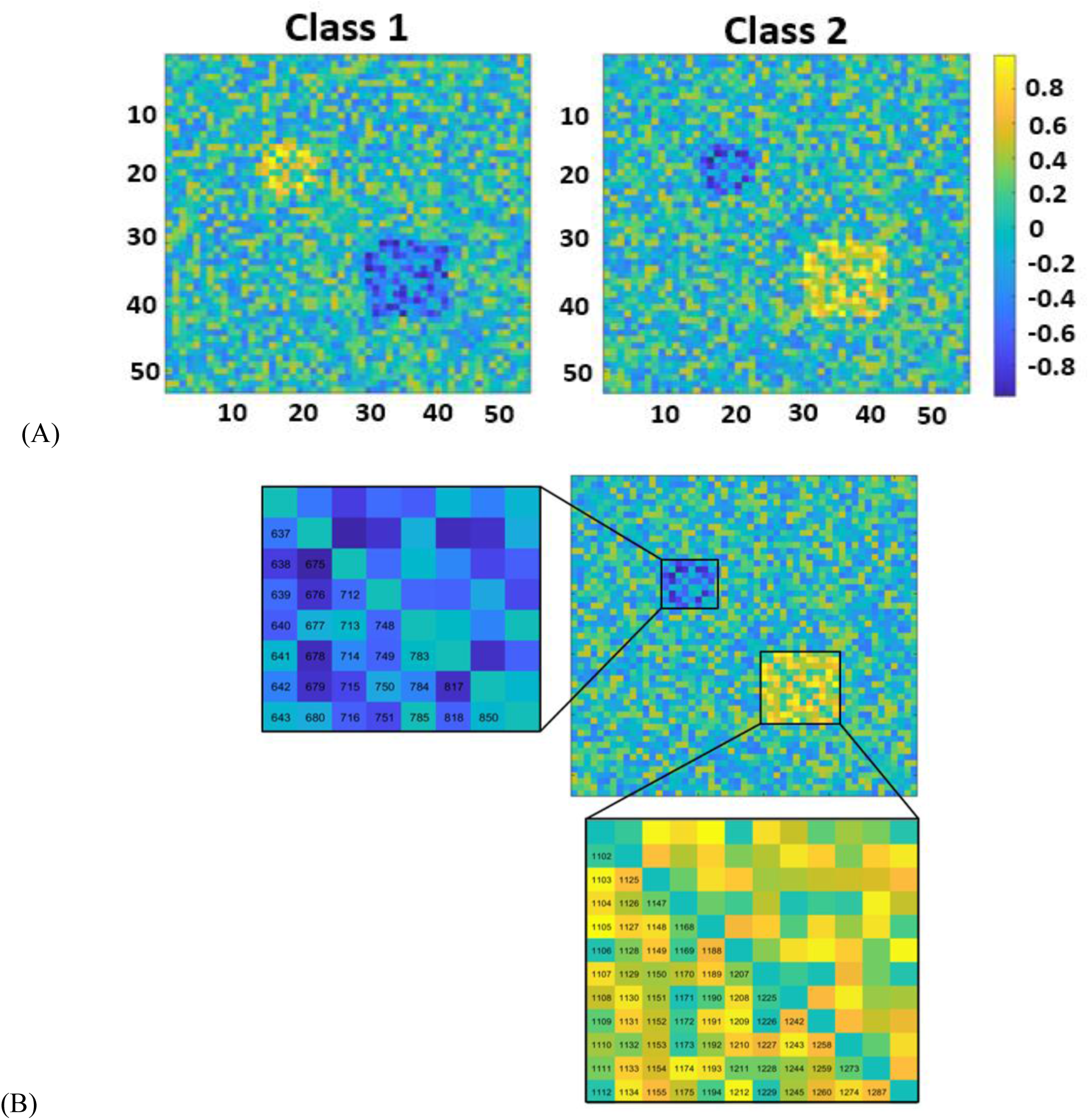
Functional Network Connectivity (FNC) in the synthetic dataset. A) The average FNC across all samples of Class 1 and Class 2. The color bar indicates the intensity of the correlation. 53×53 is the size of the connectivity matrix and hence in total we have 1378 connectivity features for each sample. We randomly generated 1378 features for each sample of both the classes. Thus, in totality we had 151 and 160 samples to mimic SZ and HC respectively (like the FBIRN dataset respectively). Then we increased the value of some of the features for one class and decreased those values for another class. B) A zoomed version of the difference between the two classes. The number is assigned to each feature as shown.

### 2.3 Classification

To classify the subjects into two classes, we trained a decision tree-based classifiers including random forest (RF), XGBoost (XGB) and CATBoost (CAT) based on the FNC features.

#### Model 1

RF is one of the most popular ensemble tree-based learning algorithms that randomly selects a subset of the training data. Then, it collects the vote from a different decision tree to assign a class to the test data. Most of the hyperparameters used in this model were default values in scikit-learn.

#### Model 2

XGB is also a very popular ensemble decision tree learning algorithm that builds the trees sequentially thereby minimizing the error of the previous tree. In this method, each tree updates the residual errors by learning from its predecessors. We used the default values in scikit-learn for most of the hyperparameters.

#### Model 3

CAT is a gradient boosting algorithm that is known for its robustness and effectiveness in handling categorical features. Unlike traditional gradient boosting methods, CAT uses an efficient method for processing categorical variables, eliminating the need for manual encoding. It builds trees in a similar sequential manner, updating residuals to minimize the loss function. The hyperparameters in CAT include the learning rate, depth of trees, number of iterations, and regularization parameters. We utilized the default values for most of the hyperparameters in the scikit-learn implementation of CAT. Additionally, early stopping was employed to prevent overfitting by monitoring the validation loss during training.

#### Classifier evaluation

A 10-fold cross-validation was employed for all three models. To assess the performance of the classifier, we calculated the area under ROC (AUC), accuracy, sensitivity, specificity, and F1 score, as shown in the equations below:

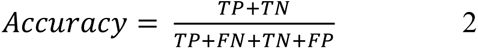

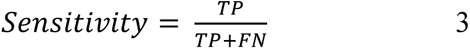

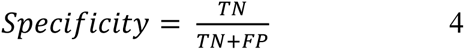

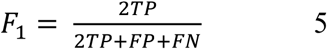

where TP, FN, TN, and FP denoted true positive, false negative, true negative, and false positive, respectively. All classifiers were implemented in Python.

### 2.4 *SHapley Additive exPlanations (*SHAP)

To explain the difference between the two classes in the trained models, we leveraged the SHAP method (https://github.com/slundberg/shap). The Shapley value estimates the magnitude and direction (or the sign) of each feature (L. S. Shapley 1953) contribution also known as a feature importance. The positive and negative signs each represent the activity and inactivity, respectively of a specific feature in the model. We computed the Shapley values for a given model, i.e., f(x), using a weighted sum that characterizes the impact of each feature being added to the model-averaged over all possible orders of features being introduced:

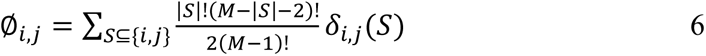

when i≠j and:

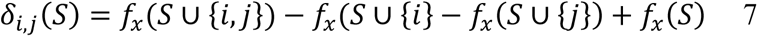

where S denotes all possible feature coalitions, *M* denotes the number of all features used in the model. The contribution or Shapley value (or ∅*i*,*j*)of each set of features, i.e., *i* and *j* here, is determined by averaging their contribution across all possible permutations of a feature set (Lundberg and Lee 2017, Lundberg *et al* 2018).

### 2.5 Dataset

This study used three datasets to validate the method. We removed the first five dummy scans before preprocessing to ensure balanced magnetization across the sequence. A smoothing kernel with an FWHM of 6 mm was applied to all datasets, as it is a widely accepted standard that minimally affects FNC and SHAP outcomes (Alaçam et al., 2023; Chen et al., 2018; Liu et al., 2017). Slice-timing correction, realignment, and smoothing were consistently applied across all datasets to maintain uniformity in preprocessing and robust SHAP feature selection.

#### 2.5.1 Synthetic Data

To test the reliability of the SHAP method, we generated synthetic data with the same number of features and samples as the real fMRI data. We first randomly generated 1378 features for each instance of both classes. In total, we generated 151 samples (to mimic SZ) and 160 samples to emulate HC groups. Then, we increased the value of some features for one class and decreased those values for another class. Therefore, two blocks of features are significantly different between the two classes (see figure 4).

#### 2.5.2 Schizophrenia (FBIRN)

The first dataset is from the Functional Imaging Biomedical Informatics Research Network (FBIRN) (T. G. M. van Erp *et al 2015*) projects. The FBIRN dataset includes seven sites containing 151 SZ subjects and 160 HC. The SZ group contains 115 males and 36 females. 36.76 is the age mean while the standard deviation is 11.63. In the HC group, we have 115 males and 35 females; the mean and the standard deviation of the age is 37.03 and 10.86, respectively. A two-sample Kolmogorov-Smirnov test was used to show that the age and sex difference between HC and SZ groups is not significant. Institutional review boards approved the consent process of each study site (S. M. Hare *et al. 2017*). The raw imaging data were collected from seven sites, including the University of California, Irvine; the University of California, Los Angeles; the University of California, San Francisco; Duke University/the University of North Carolina at Chapel Hill; the University of New Mexico; the University of Iowa; and the University of Minnesota. Imaging data were collected at six of the seven sites using a 3T Siemens Tim Trio System and at one site using a 3T General Electric Discovery MR750 scanner. Resting-state fMRI scans were acquired using a standard gradient-echo echo-planar imaging paradigm: FOV of 220 × 220 mm (64 × 64 matrices), TR = 2 s, TE = 30 ms, FA = 770, 162 volumes, 32 sequential ascending axial slices of 4 mm thickness and 1 mm skip. Subjects had their eyes closed during the resting state scan. All patients were on a stable dose of antipsychotic medication either typical, atypical, or a combination for at least 2 months. All SZs were clinically stable at the time of scanning. A diagnosis of schizophrenia is confirmed with the SCID-IV interview, and an absence of schizophrenia diagnosis in HC is confirmed with the SCID-I/NP interview. In addition, HC with a first-degree relative with an Axis-I psychotic disorder diagnosis were also excluded. As stipulated by protocol, subject symptoms were quantified using positive and negative syndrome scale (PANSS) within 1 month of imaging acquisition (Hare *et al* 2017). To ensure that demographic factors did not confound our analysis, we applied regression to remove the effects of age, sex, and imaging site from the dataset for both training and test samples within each fold separately to avoid data leakage. This step was necessary to control for the influence of age and sex on brain structure and function, as well as to account for variability across imaging sites, which can introduce unwanted variability into the dataset and reduce statistical power. By regressing out these factors, we aimed to minimize noise, increase statistical power, and enhance the interpretability of our results. Consequently, we provided a more robust analysis of resting-state functional MRI data in SZ subjects and HC.

#### 2.5.3 UK Biobank data

The data is from 9394 healthy adult individuals (average age: 63; range: 45-81 years; 4783/4611: female/male) with European ancestry available in the UK Biobank (A. U. K. Biobank 2013, C. Bycroft *et al. 2018, C. Sudlow et al. 2015*). We used a median split to put all subjects into either of the two categories: (i) old or OA (>63) and (ii) middle adult or MA (<63) groups. The middle adult group includes 4428 subjects (2406/2022: female/male), and the old group includes 4966 subjects (2377/2589: female/male). The mean age of the MA and OA groups is 55.96± 4.23 and 68.45± 3.66, respectively. The neuroimaging data was acquired on a standard Siemens Skyra 3T with a standard Siemens 32-channel RF receiver head coil. High resolution T2*-weighted functional images were acquired using a gradient-echo EPI sequence with TE =39 ms, TR = 0.735 s, flip angle = 52°, slice thickness = 3.5 mm, slice gap = 1.05 mm, field of view: 88×88×64 matrix s, voxel size = 2.4 mm 2.4 mm 2.4 mm, and 6:00 min. Written informed consent was obtained from all participants for both the UK Biobank as well as FBIRN datasets. The consent process of each study site was approved by the institutional review boards. The average FNCs across all HC (Healthy Control) and SZ (Schizophrenia) subjects in the FBRIN and Middle Adult (MA) and Old Adult (OA) in the UK Biobank are shown in Fig. 3A and Fig. 3B, respectively. In this figure the hot and cold color represent positive and negative connectivity, respectively. In order to ensure that any observed differences in functional network connectivity (FNC) were not influenced by variations in sex and imaging site, which could potentially confound the analysis results, we performed regression analysis to remove the effects of sex and site from the dataset for both training and test samples within each fold separately to avoid data leakage. This preprocessing step helped to mitigate the impact of these variables on the FNC measures, allowing us to more accurately assess the differences between groups and draw meaningful conclusions from the analysis.

### 2.6 Hyperparameter tuning of the three models for the three datasets (FBIRN, Synthetic, UK Biobank)

#### 2.6.1 Synthetic dataset

We had two classes - Class1 and Class2 and trained an RF, XGB, and CAT to classify Class1 from Class2 of the synthetic dataset. In this model, three hyperparameters were optimized via internal cross-validation, including the maximum depth level, the minimum sample split, and the minimum sample points at each node. Adam, with a learning rate of 0.001, was used for optimization. We used the default values in scikit-learn for the other hyperparameters.

#### 2.6.2 FBIRN

To classify SZ and HC subjects, we trained three tree-based classifiers: Random Forest (RF), XGBoost (XGB), and CatBoost (CAT). For the Random Forest model, we used 200 trees and optimized three hyperparameters via internal cross-validation: a maximum depth level of 90, a minimum sample split of 5, and a minimum of 2 sample points at each node. For XGBoost, we set lambda to 0.01 and alpha to 0.0001. The CatBoost model was trained with a depth of 10 and a learning rate of 0.03. We performed RandomizedSearchCV for XGBoost and GridSearchCV for CatBoost with 5-fold cross-validation to find the best parameters for each model. These best models, along with the RF model, were then fitted using 10-fold cross-validation. We iterated this process 300 times. The default values in scikit-learn were used for the other hyperparameters. The results from this 10-fold cross-validation are the basis for the SHAP graphs, AUC, and other performance metrics reported.

#### 2.6.3 UK Biobank

To classify middle-aged adults from older adults, we trained three tree-based classifiers: Random Forest (RF), XGBoost (XGB), and CatBoost (CAT). For the Random Forest model, we used 200 trees with a maximum depth level of 90, a minimum sample split of 5, and a minimum of 2 sample points at each node. For XGBoost, we set lambda to 0.01 and alpha to 0.0001. The CatBoost model was trained with a depth of 4 and a learning rate of 0.1. We performed RandomizedSearchCV for XGBoost and GridSearchCV for CatBoost with 5-fold cross-validation to find the best parameters for each model. These best models, along with the RF model, were then fitted using 10-fold cross-validation. We iterated this process 300 times. The default values in scikit-learn were used for the other hyperparameters. The results from this 10-fold cross-validation are the basis for the SHAP graphs, AUC, and other performance metrics reported.

## 3. Results

### 3.1 SHAP results on the synthetic data

We trained three classification models - RF, XGB, and CAT, based on the connectivity features from the synthetic dataset and applied the SHAP method to the model. **Figure 5** shows the top 20 features selected by the SHAP method in the synthetic dataset. Class 1 and Class 2 are represented by higher and lower SHAP values, respectively. For example, on increasing feature 1225, put the RF classifier output at Class 1, and decreasing this feature, put the RF classifier output at Class2. The increasing and decreasing patterns are akin to what we see in **Figure 3**. These results verify that the SHAP method can completely capture the difference between the two classes. We found that two out of the top 20 features overlap among all three models, as shown by purple in **Figure 5**.

**Figure 5:**
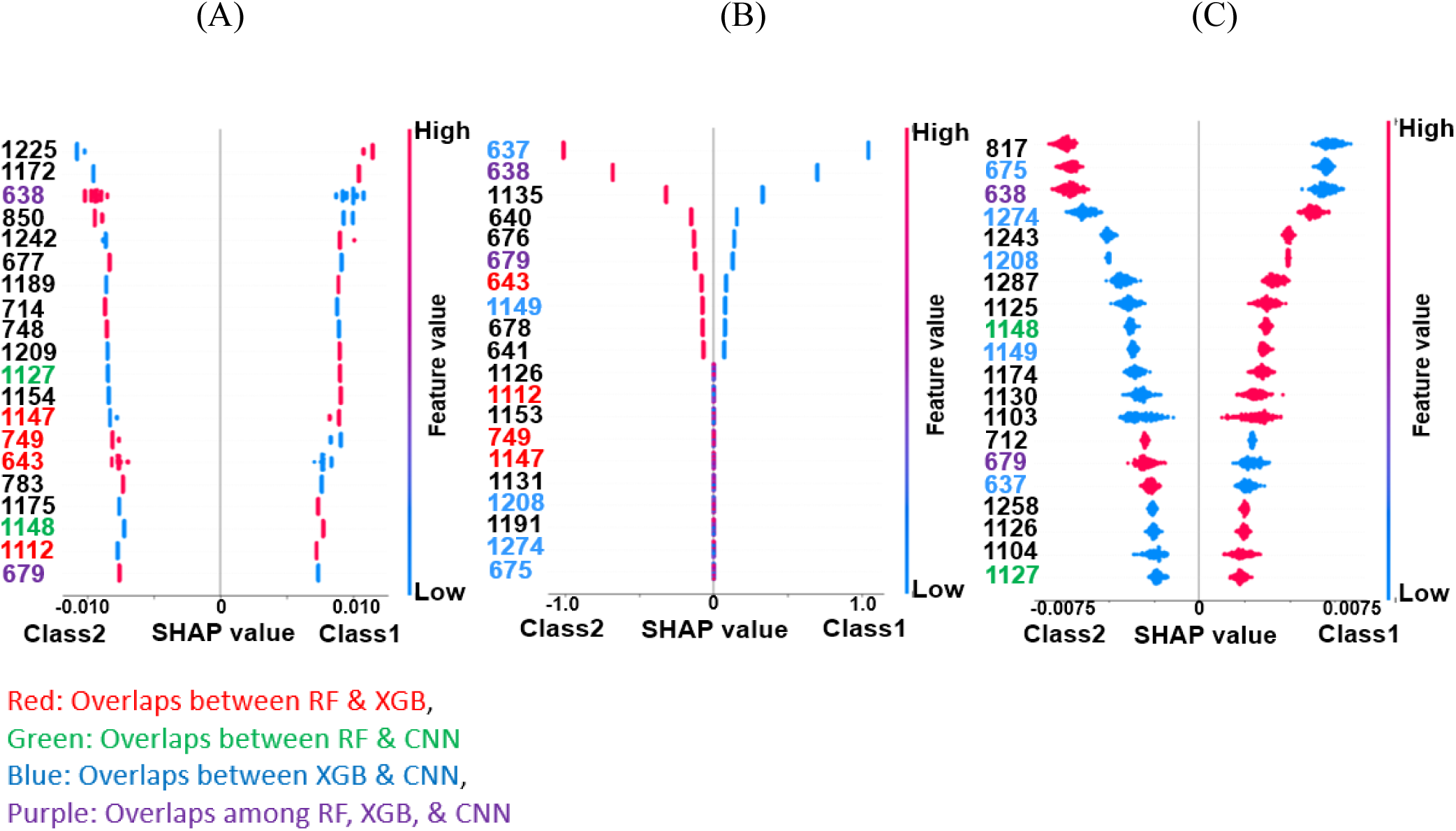
SHAP feature selection results in synthetic dataset: A) Top 20 connectivity features (out of 1378 connectivity features) of RF model selected by SHAP method. B) Top 20 connectivity features of XGB selected by SHAP method. C) Top 20 connectivity features selected by the SHAP method in CAT classifier. Also, in all graphs, the light blue shows decreasing the connectivity feature and red shows increasing the connectivity features. For example, the first connectivity features selected by SHAP method in Random Forest is feature #1225 in which increasing (red) these connectivity features would increase the likelihood of Class1 at the output of RF and decreasing (light blue) this connectivity would increase the likelihood of Class2 at the output of RF classifier.

### 3.2 SHAP results on FBIRN dataset

#### 3.2.1 Classification results between SZ and HC subject in the FBIRN dataset

Figure 6A shows the classifier receiver operating characteristic (ROCs) for RF (green), XGB (blue), and CAT (red). In addition, Table 2 shows mean accuracy, mean F1, mean sensitivity, mean specificity of these classifiers. XGB significantly outperformed the other two classifiers based on their 10-fold classification results (corrected p<0.05).

**Figure 6:**
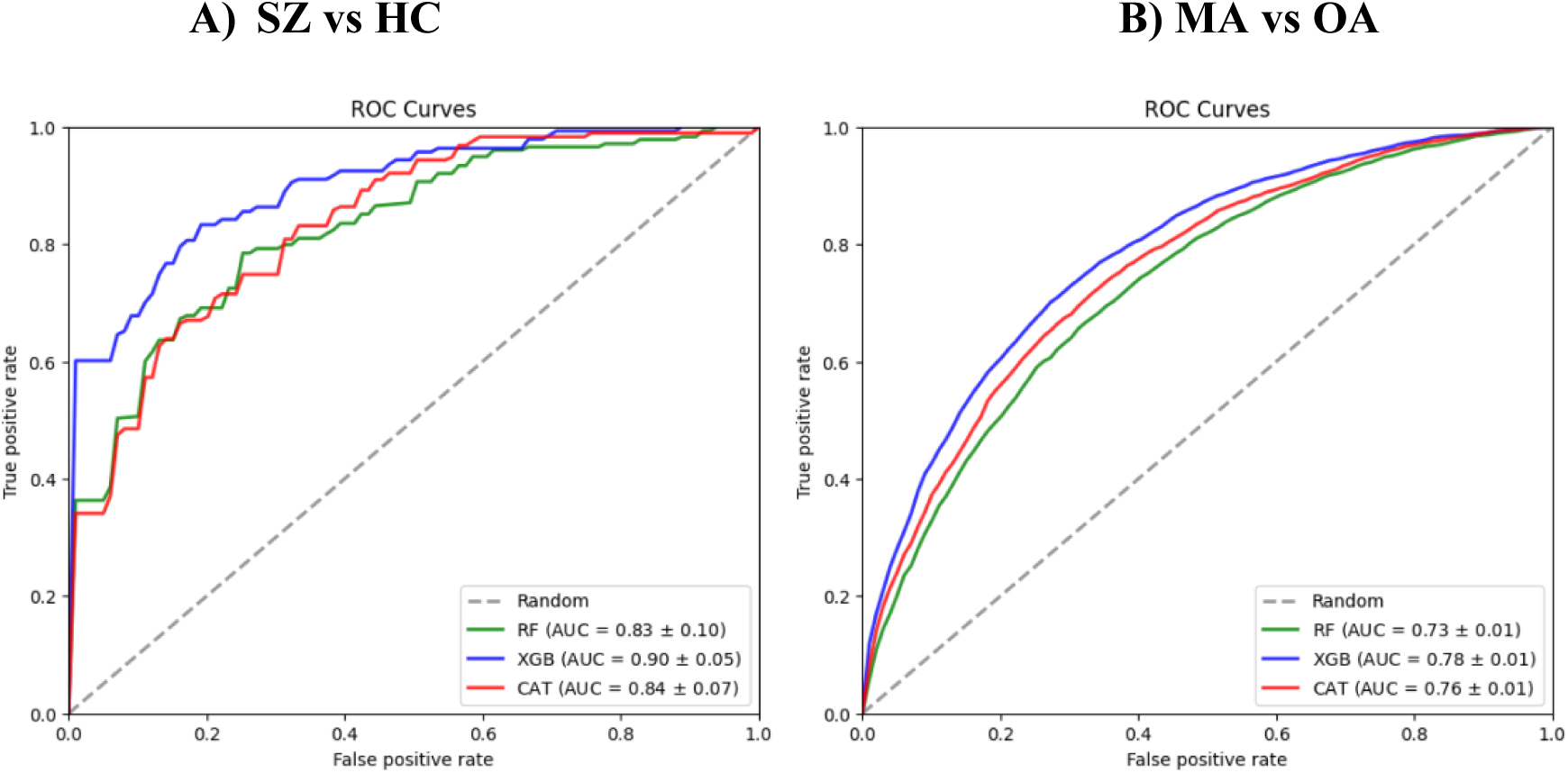
The receiver operating characteristic (ROC) curve. A) The ROC of all the classifiers that were trained in this study to differentiate between SZ and HC subjects in the FBIRN dataset. B) The ROC of all the classifiers that were trained in this study to differentiate between MA and OA subjects in the UK Biobank dataset.

**Table 2:** Performance of different classifiers in SZ/HC classification with 10-fold cross-validation.

|  | **Mean Accuracy** | **Mean AUC** | **Mean F1** | **Mean Sensitivity** | **Mean Specificity** |
| --- | --- | --- | --- | --- | --- |
| **RF** | $0.74 \pm 0.06$ | $0.83 \pm 0.03$ | $0.75 \pm 0.06$ | $0.76 \pm 0.09$ | $0.73 \pm 0.13$ |
| **XGB** | **$0.81 \pm 0.05$** | $0.90 \pm 0.02$ | $0.82 \pm 0.04$ | $0.80 \pm 0.1$ | $0.83 \pm 0.1$ |
| **CAT** | $0.72 \pm 0.07$ | $0.84 \pm 0.04$ | $0.74 \pm 0.07$ | $0.73 \pm 0.1$ | $0.72 \pm 0.14$ |

#### 3.2.2 SHAP result in the classification between SZ and HC subjects

The main focus of the current study is understanding the significance of the difference between SZ and HC subjects using an explainable machine learning approach. Using the SHAP method we found a subset of connectivity features that have the most contribution in RF, XGB and CAT model. Figure 7 shows the top 20 connectivity features in descending order that contributed more than the other features in the classification between SZ and HC subjects (for all three modes). The red and blue color in these graphs show an increase and decrease in connectivity, respectively. The positive and negative Shapley value corresponds to SZ and HC groups, respectively. In these three graphs, connectivity features that are selected only by RF and XGB are shown in red, whereas connectivity features overlapped between RF and CAT are shown in orange while those selected in both XGB and CAT are shown in blue. Finally, connectivity features that are selected by SHAP in all three models are shown in purple.

**Figure 7:**
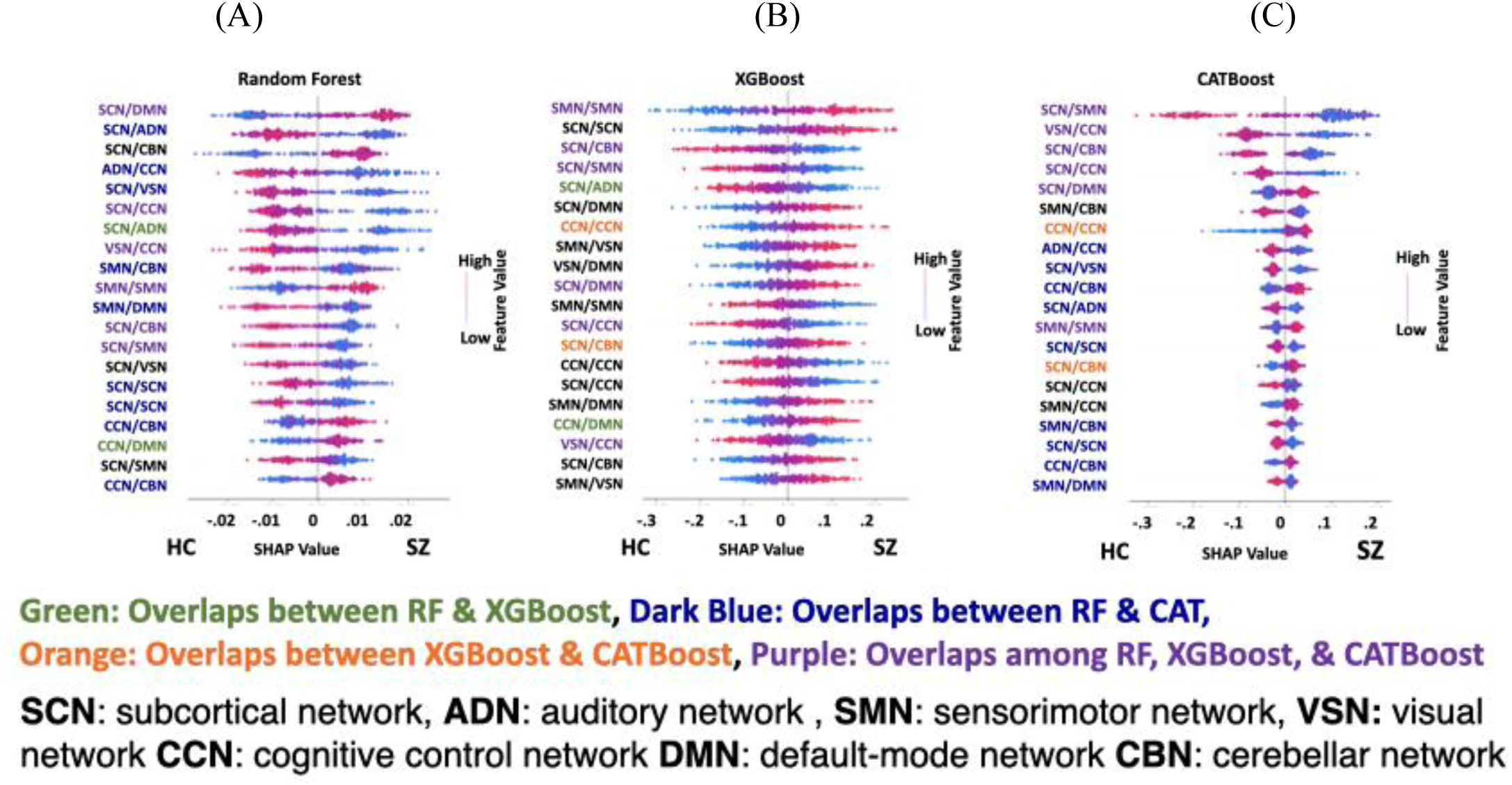
SHAP feature selection results in FBIRN dataset: **A)** Top 20 connectivity features (out of 1378 connectivity features) of RF model selected by SHAP method. **B)** Top 20 connectivity features of XGB selected by SHAP method. **C)** Top 20 connectivity features selected by SHAP method in CAT classifier. In these graphs, the connectivity feature shows the connectivity between any pair of independent components (ICAs) from subcortical network (SCN), auditory network (ADN), sensorimotor network (SMN), visual sensory network (VSN), cognitive control network (CCN), default mode network (DMN), and cerebellar network (CBN). Those connectivity features had an overlap in the feature learning results are shown in different colors. Also, all graphs light blue shows decreasing the connectivity feature and red shows increasing the connectivity features.

**Figure 7A** shows that the subcortical network (SCN) contributed the most i.e 12 out of the top 20 features in the RF model, followed by the CCN. Also, we observed that all brain networks contributed to the top 20 features selected by the SHAP in this model. A disrupted pattern was observed in the connectivity between the SCN and other brain networks. For 10 (of 12 SCN features) connectivity features related to the SCN, the connectivity increased as the likelihood of the HC group in the output of the classifier increased. Whereas for 2 (of 12 SCN features) as the connectivity related to SCN increased, the classifier’s output was the SZ group. Overall, we observed a disrupted pattern in the top 20 connectivity features selected by SHAP in the RF model.

**Figure 7B** shows the top 20 connectivity features selected by SHAP in XGB in the classification between HC and SZ subjects. Like RF, we observed that the SCN has the most contribution in the top 20 features selected by SHAP, followed by the SMN. Also, we found a disrupted (i.e., both increase and decrease) pattern in SCN connectivity in HC vs. SZ subjects. Similar to RF, we observed a disrupted pattern in the connectivity of the top 20 features in this model.

**Figure 7C** shows the top 20 connectivity selected by SHAP in the CAT model. Again, similar to the other two models, we found that the SCN contributed the most, followed by CCN just like the RF model. Similar patterns as the RF and XGB were observed in this model. We also observed a disrupted pattern in the connectivity related to SCN similar to RF and XGB.

These three graphs have 6 out of 20 features that overlapped during feature selection by SHAP in RF, XGB, and CAT. Also, we found that 2, 9 and 2 features overlapped in the model of RF and XGB, RF and CAT, and XGB and CAT, respectively.

**Figure. 8A, B, & C** visualizes the connectivity difference between HC and SZ subjects selected by SHAP in RF, XGB, and CCN, respectively. Each line represents the connectivity between a pair of components. Blue and red lines show higher connectivity in HC and SZ, respectively. All networks contribute to the top 20 features selected by the SHAP method. Also, CCN and SCN have a higher contribution in all three models. Also, we observed both an increase and decrease in the difference between SZ and HC, which proved a disrupted pattern in brain connectivity in SZ.

**Figure 8:**
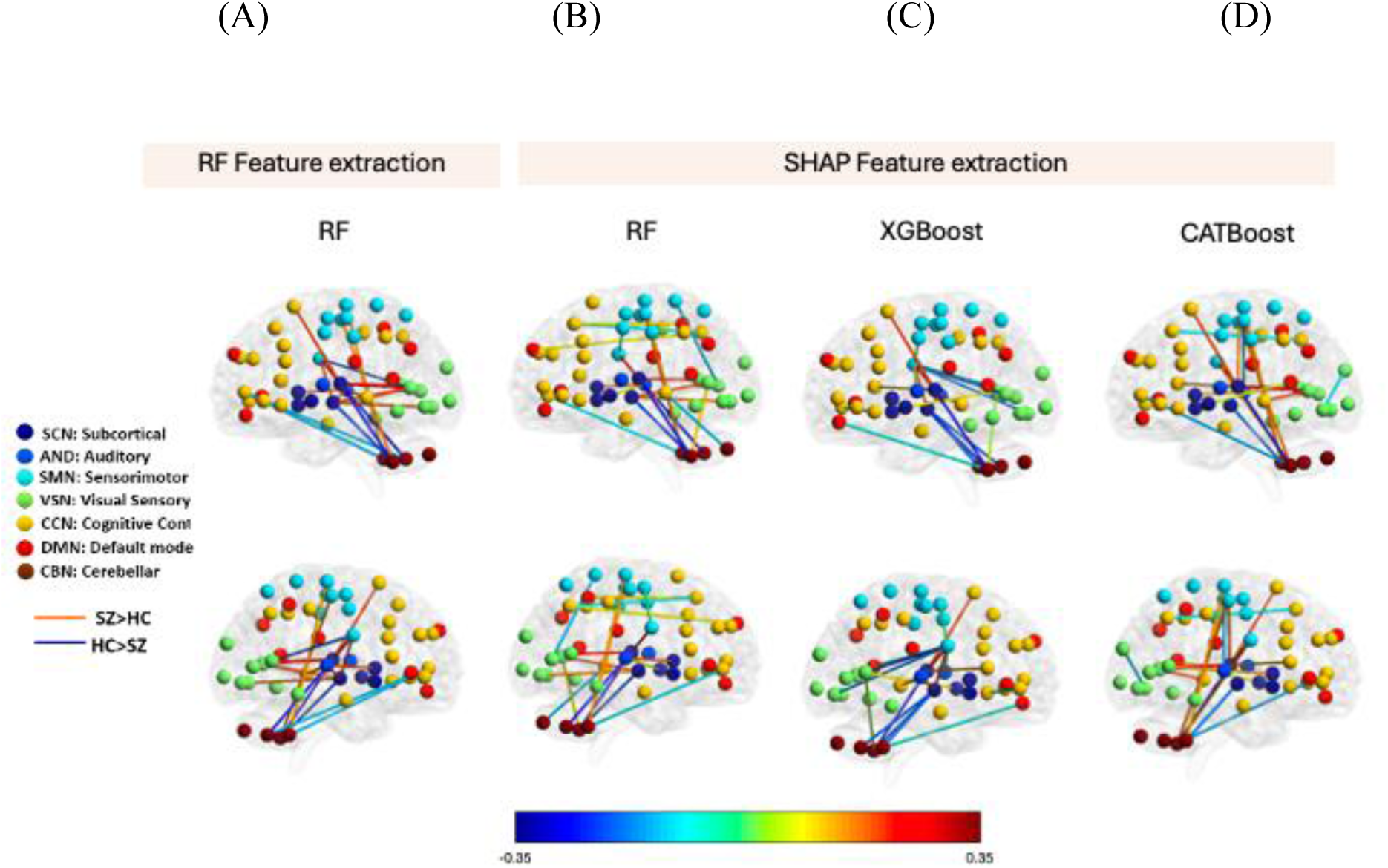
Visualization of top 20 features selected by SHAP in three models used in FBIRN dataset. We have compared our SHAP results with RF feature extraction i.e., the left-most column represents the same. **A)** Features selected by RF feature selection. **B)** Features selected by RF feature selection. SHAP in RF model. **C)** Features selected by SHAP in XGB model. **D)** Feature selected by SHAP in CAT. Each line represents the connectivity between a pair of components. Blue line shows the higher connectivity in HC and red shows the higher connectivity in the SZ. All networks contribute to the top 20 features selected by the SHAP method. Also, in all three models CCN and SCN have the higher contribution. Also, we observed both increase and decrease in the difference between SZ and HC, which proved a disrupted pattern in brain connectivity in SZ.

### 3.3 SHAP results on the UK Biobank data

#### 3.3.1 Classification results between MA and OA subject in the UK Biobank dataset

**Figure 6B** shows the classifier ROCs for RF (green), XGB (blue), and CAT (red) in the classification between MA and OA in the UK Biobank dataset. **Table 3** shows the mean accuracy, mean F1, mean sensitivity, and mean specificity of these classifiers. Overall, we observed XGB outperformed the other models.

**Table 3:** Performance of different classifiers in MA/OA classification with 10-fold cross-validation.

|  | **Mean Accuracy** | **Mean AUC** | **Mean F1** | **Mean Sensitivity** | **Mean Specificity** |
| --- | --- | --- | --- | --- | --- |
| **RF** | $0.67 \pm 0.01$ | $0.73 \pm 0.02$ | $0.68 \pm 0.01$ | $0.66 \pm 0.01$ | $0.68 \pm 0.02$ |
| **XGB** | **$0.71 \pm 0.01$** | $0.78 \pm 0.03$ | $0.71 \pm 0.02$ | $0.72 \pm 0.02$ | $0.71 \pm 0.02$ |
| **CAT** | $0.69 \pm 0.01$ | $0.76 \pm 0.02$ | $0.69 \pm 0.01$ | $0.69 \pm 0.02$ | $0.69 \pm 0.02$ |

#### 3.3.2 SHAP result in the classification between MA and OA subjects

**Figure 9A** shows the top 20 connectivity features which contributed most to the classification between MA and OA in the RF model. As shown, SCN connectivity with the rest of the brain contributed to 12 connectivity features out of the top 20 features that were selected by SHAP for classifying between OA and MA. CCN contributed the second-most to the top 20 features (i.e., 10 features) selected by the SHAP.

**Figure 9:**
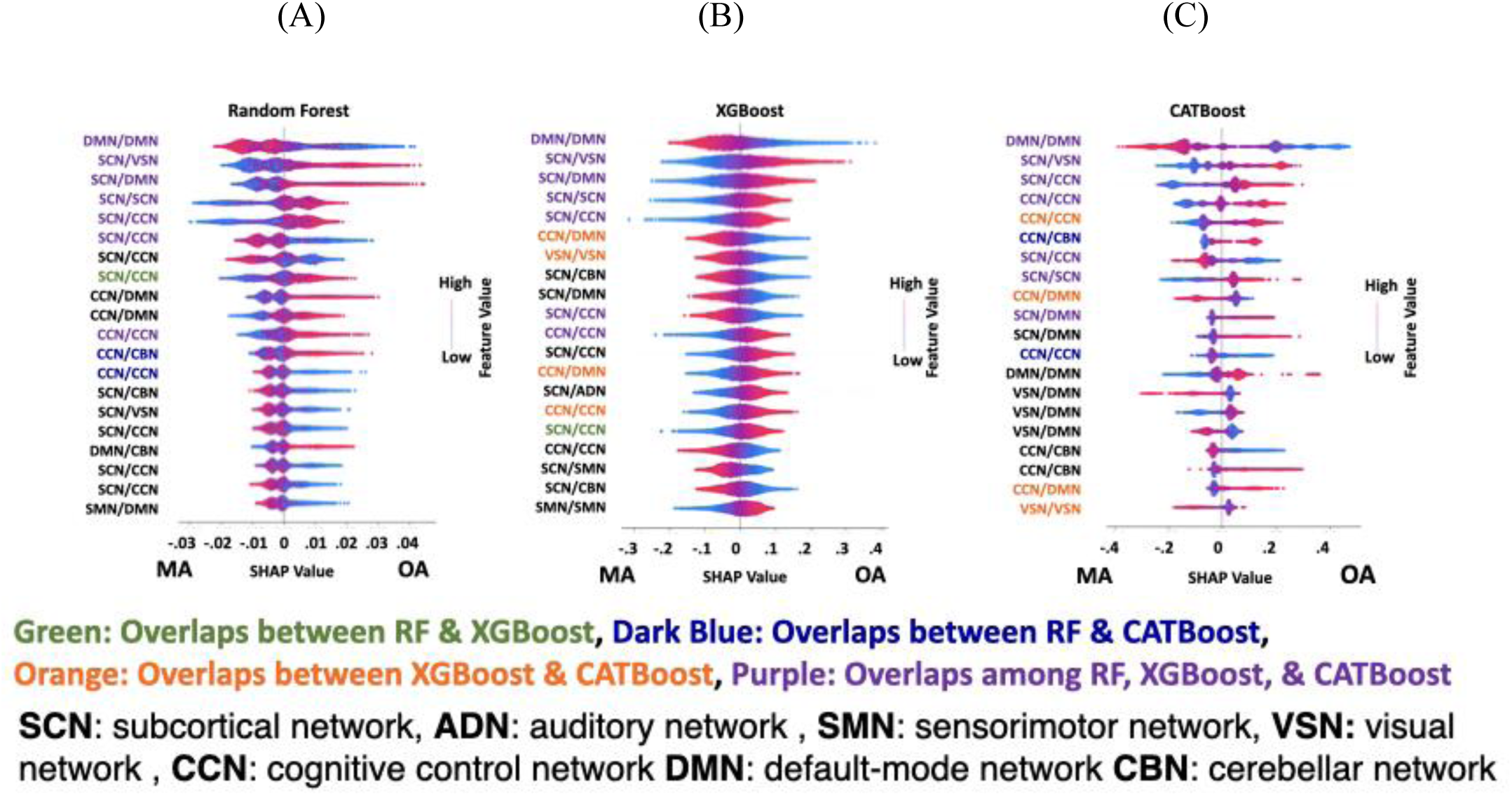
SHAP feature selection results in the classification between MA and OA in UK Biobank dataset. **A)** Top 20 connectivity features (out of 1378 connectivity features) of RF model selected by SHAP method. **B)** Top 20 connectivity features of XGB selected by SHAP method. **C)** Top 20 connectivity features selected by SHAP method in CAT classifier. Those connectivity features had an overlap in the feature learning results are shown in different color. Those features are only selected in both RF and XGB by SHAP is shown in green. Those features are only selected in both RF and CAT by SHAP is shown in dark blue. Those features are only selected in both XGB and CAT by SHAP is shown in orange. And those features are selected by SHAP method in all three methods are shown in purple. Also, all graphs light blue shows decreasing the connectivity feature and red shows increasing the connectivity features of RF classifier.

**Figure 9B** and **Figure 9C** showed that the top 20 connectivity features were selected by the SHAP method in the XGB and CAT models, respectively. In both models, SCN and CCN had the most contribution in classifying OA and MA. In addition, those connectivity features selected by the SHAP method in all three models are shown in purple. Those features only in RF and XGB are shown in green. Also, those connectivity features that are only selected in RF and CAT, in XGB and CAT, are shown in dark blue and orange, respectively.

These three graphs have 7 out of 20 features that overlapped during feature selection by SHAP in RF, XGB, and CAT. Also, we found that 1, 2 and 4 features overlapped in the model of RF and XGB, RF and CAT, and XGB and CAT, respectively.

The connectivity difference between MA and OA subjects selected by SHAP in RF, XGB, and CAT, respectively is visualized in **Figure 10**. Each line represents the connectivity between a pair of components. Blue and red lines show higher connectivity in MA and OA, respectively. All networks contribute to the top 20 features selected by the SHAP method. CCN and SCN have a higher contribution in all three models. We observed both an increase and decrease in the difference between MA and OA, which points to a disrupted pattern in brain connectivity in aging.

**Figure 10:**
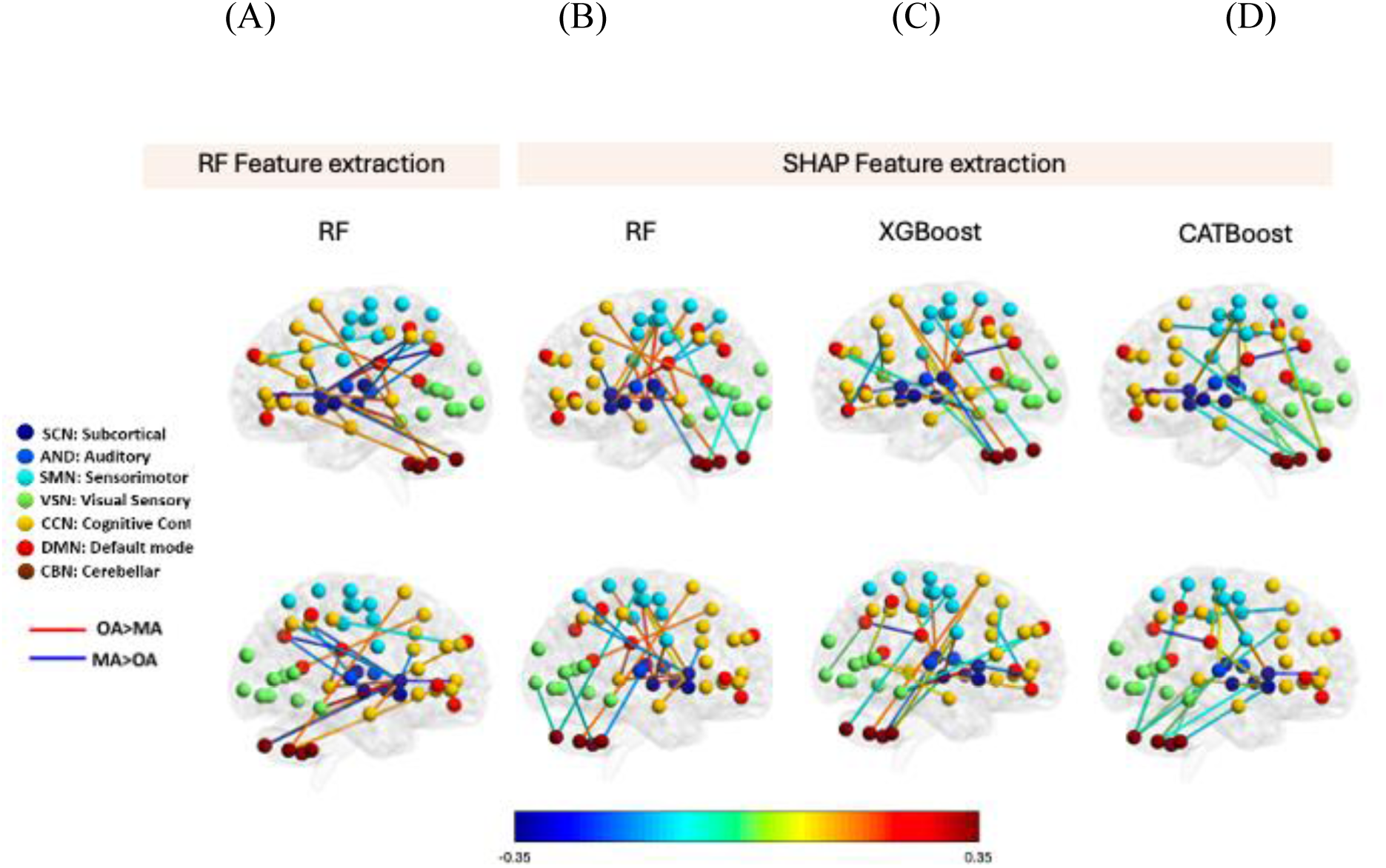
Visualization of top 20 features selected by SHAP in three models used in UK Biobank. We have compared our SHAP results with RF feature extraction i.e., the left-most column represents the same. A) Features selected by RF feature selection. B) Features selected by RF feature selection. SHAP in RF model. C) Features selected by SHAP in XGB model. D) Feature selected by SHAP in CATBoost. Each line represents the connectivity between a pair of components. Blue line shows the higher connectivity in young subjects and the red shows the higher connectivity in the old subjects. All networks contribute to the top 20 features selected by the SHAP method. Also, in all three models CCN and SCN have the higher contribution. Also, we observed both increase and decrease in the difference between middle adults and old adults, which proved a disrupted pattern in brain connectivity by progression from middle adults to old subjects.

## 4. Discussion

Our study developed a pipeline to identify the FNC features from differentiating two groups of the datasets. We validated the proposed framework on three models with the help of the synthetic dataset. This framework was validated on a synthetic dataset and two datasets, including FBIRN (T. G. M. van Erp *et al 2015*) and UK Biobank dataset (A. U. K. Biobank 2013, C. Bycroft *et al. 2018, C. Sudlow et al. 2015*). In all models, our framework successfully selected only the specific features that varied between two classes of the synthetic data.

In this study, we focused on validating the effectiveness of our pipeline rather than comparing its performance with previous models. By applying it to both synthetic and multiple real datasets across different models, we demonstrated its robustness and generalizability in various settings. This comprehensive validation highlights the reliability of our approach and underscores its contributions to the existing body of knowledge. Our findings provide a solid framework for further research and practical applications, offering a dependable methodology that can be adopted in diverse contexts within the field.

### 4.1 Schizophrenia biomarker

The current study explored the functional connectivity among 7 data-driven networks, which included SCN, ADN, VSN, SMN, CCN, DMN, and CBN, to differentiate individuals with schizophrenia (SZ) from control (HC). We trained three models, including RF, XGB, and CAT and found that all three models could successfully differentiate between HC and SZ. XGB showed the highest classification accuracy of 81.03%. We then estimated the top 20 features out of 1378 which had the most significant contribution for classifying SZ and HC. To summarize the connectivity pattern between SCN and SMN, VSN and CCN, SCN and ADN were constant across the different model architectures i.e., RF, XGB and CAT. In all the three pairs, increasing the connectivity strength, increased the likelihood of HC group as the output of the model. Thus, found that 6 of the top 20 selected features overlap across all three models. 2 features overlapped between RF and XGB, 9 between RF and CAT, and 2 between XGB and CAT. Regarding the interpretability of the SHAP method, we conducted additional analysis to support our claims. For the SZ dataset, the RF model showed an 85% overlap with at least one other model when considering the top 20 features, indicating high consistency in feature importance. The XGBoost model had a 50% overlap, and CATBoost demonstrated an 85% overlap with at least one other model, all based on the top 20 features. Notably, there was a 30% overlap among all three models, underscoring the robustness of the identified features. This amount of overlap on the top 20 features selected by SHAP in three different models is a good testament to the robustness of the proposed framework.

We found that all seven networks significantly contributed to discrimination between SZ and HC. This finding aligns with previous literature, which has highlighted the widespread and varied nature of network dysfunction in schizophrenia (Garrity et al., 2007; Cole et al., 2013; Salvador et al., 2010). We also found contribution of SMN from our analysis in SZ which has also been mentioned in literature (Hummer et al 2020). Additionally, studies have shown that schizophrenia affects multiple brain networks, encompassing sensory, motor, and cognitive domains (Kaufmann et al., 2015; Berman et al., 2016; Ray et al., 2017). Given the pervasive nature of these disruptions, our approach of considering contributions from all seven networks helps address the challenge of false positives. By leveraging a comprehensive analysis across diverse networks, we ensure that the classification is not driven by isolated anomalies but by consistent patterns of connectivity disruptions across the brain. This methodology is supported by recent findings that highlight the regional, circuit, and network heterogeneity of brain abnormalities in psychiatric disorders (Segal et al., 2023). In contrast to the main body of previous research, which only focused on the functional connectivity of DMN in schizophrenia (W. Guo *et al. 2017*), we found that functional connectivity of other networks, in particular CCN and SCN, also contributes significantly to the discrimination between SZs and HCs. Our findings regarding CCN are coherent with previous literature (Horne, M et al. 2021; Culbert A, et al. 2015, Alústiza, I et al. 2016). We also found some literature (Andreasen N, et al. 1998; Salvador et al 2010; Zhang et al 2012) mentioning the contribution of SCN in schizophrenia which aligns with our findings as well. Our analysis benefits from extensive evidence that schizophrenia manifests across multiple networks, as indicated by our high overlap in feature importance across different classifiers. This widespread involvement of various networks reduces the likelihood that our results are due to random or isolated connectivity issues. Instead, it reflects the robust and consistent patterns of network dysfunction characteristic of schizophrenia (Ji et al., 2019; Lesh et al., 2011; Sendi et al., 2021). By accounting for the multifaceted nature of network disruptions, we provide a more accurate and comprehensive understanding of the neural underpinnings of schizophrenia.

### 4.2 The aging biomarker

The current study explored the functional connectivity among 7 data-driven networks, including SCN, ADN, VSN, SMN, CCN, DMN, and CBN between older adults (≥ 63) or OA and middle adults (<63) or MA subjects using rs-fMRI of the UK Biobank (UKBB) data set. We trained and tested three tree-based models-RF, XGB, and CAT. Using these models, we could differentiate OA from MA subjects with a mean accuracy of more than 67%. XGB showed the highest accuracy of 71.38%. In the UKBB dataset, the RF model showed a 50% overlap with at least one other model, while XGBoost had a 60% overlap and CATBoost demonstrated a 65% overlap with at least one other model for the top 20 features. Additionally, a 35% overlap among all three models was observed. These findings highlight the consistency of the identified connectivity features across different classifiers, providing a clearer understanding of the robustness of our results. This result highlights the contribution of the brain FNC in the classification of OA and MA.

We also found a subset of features that have the highest contribution to the classification between OA and MA subjects in each model. We discovered that 25% of the top 20 features were shared among all three models. We also found that all 7 networks contributed to all top 20 connectivity features selected by the SHAP method in all three models. This is consistent with previous studies, which showed the effect of age on the between-network connectivity in the adult subjects (L. Geerligs et al. 2015, M. Y. Chan et al. 2014). In contrast to the study mentioned above, in which the statistical learning method was used to find the difference in the FNC between the OA and MA subjects, our current study used a novel feature learning method to model this difference. In contrast to statistical learning, which typically evaluates each feature one by one and does not consider the interaction between input features, the machine learning-based feature learning approach provides a generalized model of the difference between the older adult and middle adult features (D. Bzdok et al. 2018).

By comparing the connectivity between OA and MA subjects of the top 20 features, we found a pattern of increase and decrease. That possibly showed a disrupted pattern in the whole-brain FNC, consistent with the previous literature (H. I. Zonneveld *et al. 2019*). We finally found that CCN and SCN contribute more than the other networks in this classification, and this result is consistent among all models. This finding is consistent with previous literature, where the role of CCN and SCN was highlighted (M. Ystad et al. 2010, K. L. Campbell et al. 2012, Wierenga et al. 2008, Tomasi et al. 2012).

### 4.3 Comparing SHAP with other feature selection method and existing literature

Conventional statistical learning methods suffer from low prediction accuracy, albeit being able to model the relationship between variables (Butler et al. 2015, Bass et al. 2015, Mohan et al. 2020, Bronte-Stewart H et al. 2009). Besides, as the number of input variables increase, the inference in statistical learning can become less precise. On the other hand, machine learning methods focus on prediction accuracy which utilizes the generalizable learning algorithm. It finds a pattern of high-dimensional data across patients (i.e., leave-one-patient-out) (Habets JGV et al. 2020). Although there are compelling prediction results in machine learning methods, lack of an interpretable model makes these methods useless to understand the fundamental mechanism of the current neurophysiological knowledge (Bzdok et al. 2018). Recent progress in machine learning algorithms, particularly in feature learning methods, provides various interpretability degrees, from the highly interpretable linear models such as Least Absolute Shrinkage and Selection Operator (LASSO) regression to black-box models such as neural networks (Cai J et al. 2017). Additionally, in comparison with statistical testing methods, which do not account for the joint space interactions of markers associated with neurological activity; and cannot indicate the likelihood of inter-subject generalizability of identified biomarkers; interpretable machine learning accounts for those.

SHAP has also advantages over the other interpretable approaches used. While LASSO is effective for selecting a subset of features when the number of samples is much smaller than the number of features, it may struggle when the opposite scenario arises. Additionally, LASSO is limited to specific models and may only choose one feature from a group of highly correlated features. In contrast, SHAP (SHapley Additive exPlanations) offers a model-agnostic approach to feature importance analysis by describing importance and contributions as the sum of the feature contributions to an outcome, making it applicable to a wide range of machine learning and deep learning models (Gramegna et al. 2022). At the same time, some other methods like LASSO (L1 norm) or elastic net (L1 and L2 norms) are restricted to a specific model. Additionally, LASSO for *n*<<*p* (n is the number of samples and *p* is the number of features), LASSO selects at most *n* features (M. S. E. Sendi *et al. 2021*). Additionally, LASSO only uses random selection of one feature from a group of highly correlated features. Whereas SHAP will find top features across the whole set of features and does not focus on small regions in the feature space. Also, SHAP values show the feature’s importance by indicating whether the feature has a positive or negative impact on the classifier output.

Gini impurity is another widely used metric for feature extraction, which usually is used decision tree algorithms, measuring the impurity of a dataset by evaluating the probability of misclassifying a randomly chosen element. It efficiently partitions the feature space based on the homogeneity of the target variable. However, it may not capture complex interactions between features. In contrast, SHAP provides a model-agnostic approach to interpret the output of any machine learning model. By attributing the contribution of each feature to the model’s predictions, SHAP offers insights into the importance and impact of individual features on the model’s output. Despite its interpretability and versatility, SHAP may require extensive computational resources. Nevertheless, it remains a valuable tool for understanding model behavior and making informed decisions in various domains. (Deb et al. 2021, C Müller 2017, Lundberg et al, 2017). Secondly, permutation feature importance is commonly used to gauge the significance of individual features in machine learning models, yet its efficacy can be hampered by various constraints. Perturbing single features may not notably affect model performance and executing permutation feature importance analyses can be computationally demanding, particularly with a complex model. In contrast, SHAP provides a more comprehensive and interpretable approach to feature importance analysis, making it advantageous for understanding model predictions and decision-making (Lundberg et al, 2017, Altmann et al. 2010).

SHAP also offers several advantages over Layer-wise Relevance Propagation (LRP), particularly due to its model-agnostic nature and robust theoretical foundation. Unlike LRP, which is specifically designed for neural networks, SHAP can be applied to any machine learning model, including but not limited to decision trees, ensemble methods, support vector machines, and neural networks. This versatility makes SHAP extremely valuable for comparative analysis across different models, enabling consistent interpretation regardless of the underlying algorithm. Moreover, SHAP is grounded in cooperative game theory, providing a fair and theoretically justified distribution of contributions among features based on Shapley values. This not only ensures equitable attribution but also enhances the interpretability and trustworthiness of the explanations. SHAP also excels in visualization capabilities, offering a variety of plots that can articulate the influence of each feature on model predictions in a clear and impactful way. While SHAP might be computationally intensive due to its need to evaluate all possible combinations of features, its comprehensive insights and adaptability across diverse scenarios give it a significant edge in the field of explainable artificial intelligence (Lundberg and Lee 2017).

Our study specifically targets disease classification using resting-state fMRI data, distinguishing it from the task-based fMRI focus of (Narun Pat et al. 2022), which examined the relationship between task-based fMRI and individual cognitive differences. Resting-state fMRI, unlike task-based fMRI, captures intrinsic brain activity patterns without explicit tasks, providing insights into the brain’s functional connectivity and organization. (Vedaei et al. 2023) utilized resting-state fMRI and machine learning for chronic mild traumatic brain injury (TBI) classification. Our research extends beyond a single neurological condition to classify various neurological and psychiatric disorders, incorporating advanced feature extraction and interpretable machine learning models to enhance robustness and interpretability. Additionally, unlike (Kim et al. 2022), who predicted social anxiety using resting-state fMRI, our study adopts a disease-agnostic approach, enabling the identification of common underlying patterns of brain dysfunction across multiple disorders. By employing rigorous cross-validation strategies and addressing confounding variables such as age, sex, and site effects, we ensure the generalizability and reliability of our findings. In our study, we selected the top 20 features out of 1378 connectivity features for each sample to enhance the classification between the two classes. This choice, guided by the default setting in the SHAP toolbox for Python, strikes a balance between providing detailed insights and maintaining clarity in visualization. While selecting more features could clutter the visual interpretation and selecting fewer might omit significant patterns, we opted for 20 to ensure an optimal balance. This approach is consistent with previous research, such as the works by Tasnim et al. (2023), Mi et al. (2021), and AlMansoori et al. (2024), where a similar number of features is chosen to maintain clarity and comprehensiveness in visualizations. Thus, our methodology ensures that the most impactful features are covered comprehensively, aligning with established practices in the field and contributing to a more robust and interpretable model. These distinctive features underscore our contributions to neuroimaging-based disease diagnosis and classification, with significant implications for clinical practice and translational neuroscience research.

### 4.4 Limitations and future work

This study uses a different interpretability approach and applies it to two neurological datasets and a synthetic dataset as well as uses different models (RF, XGB, CAT). Inspite of this, there are a few limitations associated with this study besides underspecification (D’Amour et al. 2020). First, we only used the SHAP method as an interpretable approach to differentiate two groups of samples based on the whole-brain FNC. Therefore, future work is needed to explore other explainable methods of which some could also include the ensemble models like VarGrad and SmoothGrad-Squared (Hooker et al. 2019) and compare their results with SHAP. To increase the reliability of the model and to make it more robust we could also include simulations with input invariance and look for consistencies (Kinderman et al 2019). Secondly, the performance of other linear and nonlinear machine learning models needs to be evaluated for classifying between two groups based on the FNC. It would be interesting to experiment in more depth using the utility metrics if there is a significant difference including and excluding the results from explainability approaches like SHAP (Weerts et al. 2019). Thirdly, we have only tested with the fMRI modality, investigating other modalities like structural MRI, Diffusion Tensor Imaging (DTI) would be interesting to see if there is consistency or association between the findings.

### 4.5 Conclusion

Our work proposed a framework to identify the FNC biomarkers through the SHAP method which differentiates two groups of samples. We validated the framework’s robustness in three datasets, which included FBRIN, UK Biobank and a synthetic dataset. While we proved that the framework finds only those FNC features that are different from two classes of the synthetic dataset, we found the FNC biomarker which is associated with schizophrenia and aging.

## Author’s contribution

Mohammad S. E. Sendi and Vaibhavi Itkyal contributed equally for this manuscript and indicates joint first authorship. Sendi and Itkyal developed the study, conducted data analysis, interpreted the results, and wrote the original manuscript draft. Sabrina conducted data analysis. Vaibhavi Itkyal wrote the original manuscript draft. Z. Fu conducted the preprocessing analysis. Vince D. Calhoun developed the study, interpreted the results, edited the original draft, secured funding, and provided critical thinking of the initial draft. Sendi and Calhoun were involved in supervising the study. All authors approved the final manuscript.

## Code and data availability statement

The code used for preprocessing and FNC calculation are available at https://trendscenter.org/software/. Also, statistical parametric mapping (SPM 12) is available at https://www.fil.ion.ucl.ac.uk/spm/.The Neuromark framework and the Neuromark template (Neuromark_fMRI_1.0) have been made available and incorporated into the Group ICA Toolbox (GIFT v4.0.5.14: https://trendscenter.org/software/gift/). Users worldwide can now directly download and utilize these resources. We also use this https://www.nitrc.org/projects/bnv/ for brim graph.

## Acknowledgements

We thank those who collected the data and the participants of this study.

## Supplementary tables

**Table 1:** UK Biobank Dataset Summary.

|  |  | Middle Aged adults (MA) | Old aged adults (OA) |
| --- | --- | --- | --- |
| Site 1 (Site == 11025) | Number | 3044 | 2786 |
| | Age | $57.44 \pm 4.47$ | $70.04 \pm 3.63$ |
|  | Gender(M/F) | 1397/1647 | 1535/1251 |
| Site 2 (Site == 11026) | Number | 517 | 672 |
| | Age | $58.29 \pm 4.08$ | $71.23 \pm 3.89$ |
|  | Gender(M/F) | 238/279 | 338/334 |
| Site 3 (Site == 11027) | Number | 1136 | 1239 |
| | Age | $58.17 \pm 4.04$ | $70.78 \pm 3.94$ |
|  | Gender(M/F) | 492/644 | 611/628 |

**Table 2:**
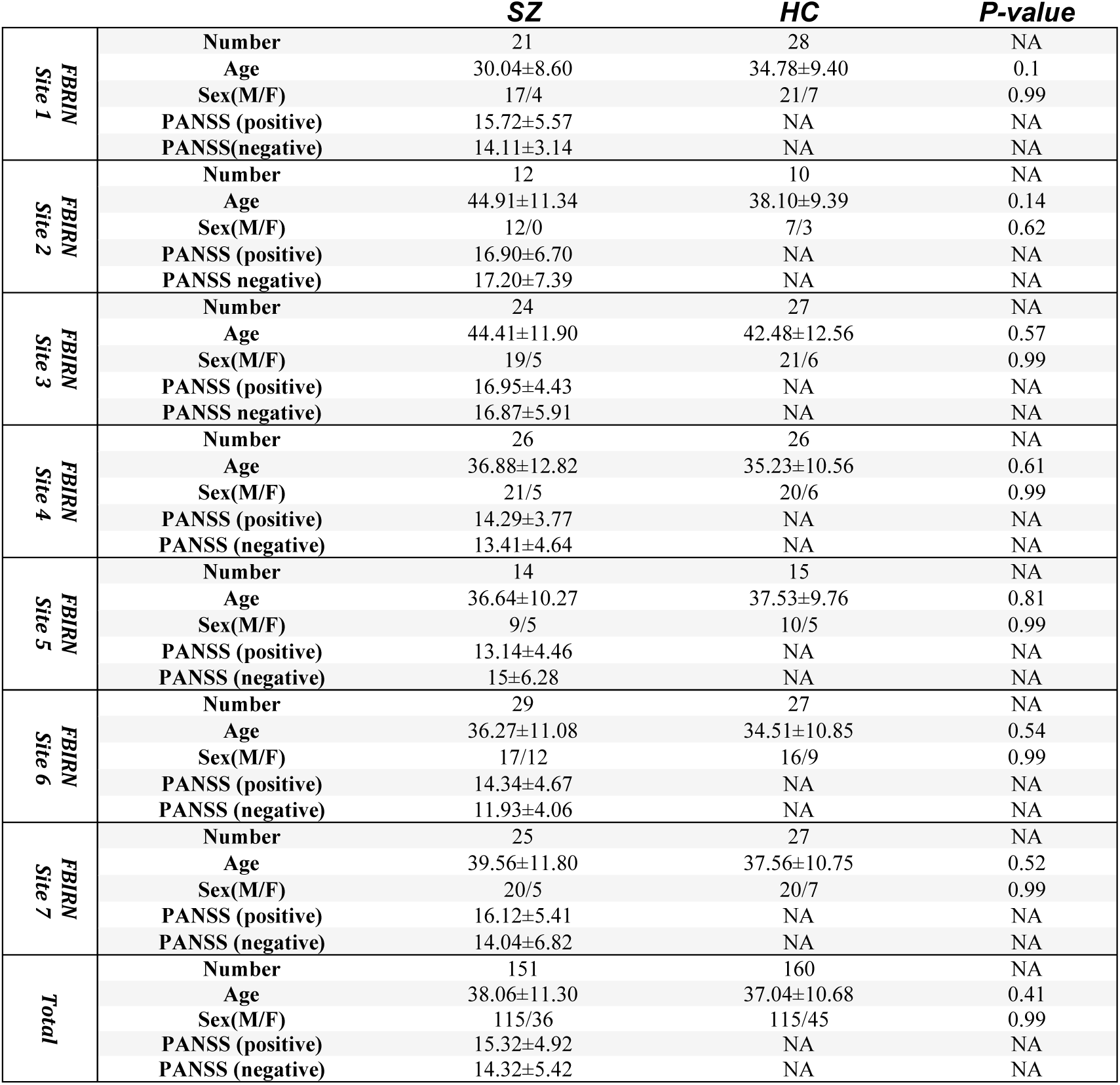
FBIRN Dataset Summary.

